# Stress- and pathway-specific impacts of impaired jasmonoyl-isoleucine (JA-Ile) catabolism on defense signaling and biotic stress resistance in Arabidopsis

**DOI:** 10.1101/686709

**Authors:** Valentin Marquis, Ekaterina Smirnova, Laure Poirier, Julie Zumsteg, Fabian Schweizer, Philippe Reymond, Thierry Heitz

## Abstract

Jasmonate (JA) synthesis and signaling are essential for plant defense upregulation upon herbivore or microbial attacks. Stress-induced accumulation of jasmonoyl-isoleucine (JA-Ile), the bioactive hormonal form triggering major transcriptional changes, is often dynamic and transient, due to the existence of potent removal mechanisms. Two distinct but interconnected JA-Ile turnover pathways have been described in Arabidopsis, either via cytochrome P450 (CYP94)-mediated oxidation, or through deconjugation by the amidohydrolases (AH) IAR3 and ILL6. Their impact was not well known because of gene redundancy and compensation mechanisms when each pathway was partially impaired. Here we address the consequences of fully blocking either or both pathways on JA homeostasis and defense signaling in three mutant backgrounds: a double *iar3 ill6* (*2ah*) mutant, a triple *cyp94b1 b3 c1* mutant (*3cyp*), and a newly generated quintuple (*5ko*) mutant deficient in all known JA-Ile-degrading activities. These lines behaved very differently in response to either mechanical wounding, insect attack or fungal infection, highlighting the stress-specific contributions and impacts of JA-Ile catabolic pathways. Deconjugation and oxidative pathways contributed additively to JA-Ile removal upon wounding, but their genetic impairement had opposite impacts on *Spodoptera littoralis* larvae feeding: *2ah* line was more resistant whereas *3cyp* was more susceptible to insect attack. In contrast, *2ah, 5ko* but not *3cyp* overaccumulated JA-Ile upon inoculation by *Botrytis cinerea*, yet *3cyp* was most resistant to the fungus. Despite of building-up unprecedented JA-Ile levels, *5ko* displayed near WT levels of resistance in both bioassays. Molecular and metabolic analysis indicated that restrained JA-Ile catabolism resulted in enhanced defense and resistance levels only if genes encoding *JAZ* or *JAM* negative regulators were not simultaneously overstimulated. Our data demonstrate that despite of acting on a shared hormonal substrate, AH or/and CYP94 deficiency differentially impacts JA homeostasis, responses and tolerance to related biotic stresses.

## INTRODUCTION

Jasmonates (JAs) have been recognized for more than two decades as powerful regulators of inducible defense responses protecting plants from damage inflicted by herbivorous insects or microbial pathogens (Campos et al. 2014; Wasternack and Hause 2013). Even before the identification of jasmonoyl-isoleucine (JA-Ile) as the critical bioactive form in 2007 (Chini et al. 2007; Thines et al. 2007), many derivatives of jasmonic acid (JA) were described, resulting from hydroxylation, conjugation to sugars (Miersch et al. 2008), amino acids or amino cyclopropane carboxylic acid, sulfation (Gidda et al. 2003), decarboxylation and many more (Wasternack and Song 2017). The enzymes responsible for these modifications have not all been identified and the function of these derivatives, if any, are generally unknown. The master regulator jasmonoyl-(L)-isoleucine (JA-Ile) results from a specific conjugation reaction of jasmonic acid (JA) by the enzyme JASMONATE RESISTANT 1 (JAR1). In the core perception and signaling pathways, JA-Ile acts as a ligand promoting the assembly of the co-receptor CORONATINE INSENSITIVE 1 (COI1) with distinct JASMONATE ZIM-DOMAIN PROTEIN (JAZ) that otherwise powerfully repress target transcription factors and responses. After assembly, COI1-JAZ is recruited into the E3 ubiquitin ligase SCF^COI1^ that tags JAZs for proteolytic degradation, and provides the basis for JA-Ile-responsive gene de-repression (Chini et al. 2007; Thines et al. 2007). Several hundreds to thousands of genes are under this control, and depending on developmental stage, organ, nature of stimulus, and crosstalks with other hormones, a wide array of JAZ-transcription factor combinations provide specificity to the system with only one major hormonal ligand (Chini et al. 2016). Distinct JA-Ile-triggered networks regulate leaf defenses against different types of aggressors and are integrated by separate sets of transcription factors (TF). MYC2, a bHLH type TF, integrates (together with MYC3 and MYC4) simultaneous JA/abscisic acid stimuli and defines a wound/insect-specific branch reflected by induction of markers such as vegetative storage protein (VSP); ERF1/ORA59 TFs integrate concomitant JA/ethylene signals into a microbe-specific branch probed by plant defensin *PDF1.2* (Pieterse et al. 2012; Wasternack and Hause 2013).

JA signaling, like any hormonal pathway, needs to be tightly controlled in time and space, particularly because defense upregulation is costly and linked to overall growth inhibition. This antagonism was long thought to be merely imposed by limited resources that need to be re-allocated from developmental to defense sinks (Havko et al. 2016), but recent advances have shown that additional hardwire connections control growth-defense tradeoffs (Campos et al. 2014; Guo et al. 2018). For appropriate control of JA responses, plants rely on a number of negative feedback mechanisms. In addition to JAZ repressor proteins, other negative regulators were identified that repress JA responses at the promoter level, including JAV (Yan et al. 2018) and the JAM subclass of bHLH (Sasaki-Sekimoto et al. 2013) proteins. Another way to terminate jasmonate action is at the metabolic level by eliminating the active ligand JA-Ile. The characterization of JA-Ile catabolic pathways has shed light on hormone homeostasis regulation and has also provided additional complexity in the JA metabolic grid. Two distinct enzymatic pathways are known to modify or degrade the JA-Ile hormone. One consists in a two step oxidation of the terminal (ω) carbon of the JA moiety by cytochrome P450 monooxygenases of the CYP94 family. In Arabidopsis, CYP94B3 and to a minor extent CYP94B1 generate 12OH-JA-Ile as a main product whereas CYP94C1 catalyzes the full oxidation of JA-Ile to 12COOH-JA-Ile (Figure 1) (Bruckhoff et al. 2016; Heitz et al. 2012; Kitaoka et al. 2011; Koo et al. 2011, 2014). The first oxidation product retains reduced receptor-assembly capacity and gene-inducing activity (Aubert et al. 2015; Koo et al. 2011; Poudel et al. 2019), but the second product proved fully inactive (Aubert et al. 2015; Koo et al. 2014). The second pathway is defined by deconjugating activity, first identified in *Nicotiana attenuata* (Woldemariam et al. 2012) and subsequently characterized in Arabidopsis as mediated by the amidohydrolases IAR3 and ILL6 (Figure 1) (Widemann et al. 2013). These enzymes act on JA-Ile and other JA-amino acid conjugates but also on 12OH-JA-Ile generated by CYP94Bs, releasing JA and 12OH-JA from these respective substrates (Widemann et al. 2013; Zhang et al. 2016). The interconnection of the two catabolic pathways generates increased metabolic complexity because many of the above-mentioned JAs derive in fact from JA-Ile catabolism (Heitz et al. 2016; Widemann et al. 2013). Both JA-Ile oxidases and amidohydrolases genes are induced by environmental cues and are co-regulated with the central regulon defined by JA pathway biosynthetic and signaling genes. Their simultaneous enzymatic action shapes specific jasmonate patterns in different tissues and stress conditions (Aubert et al. 2015; Heitz et al. 2012; Koo and Howe 2012; Koo et al. 2014; Widemann et al. 2013, 2016).

**Figure 1.**
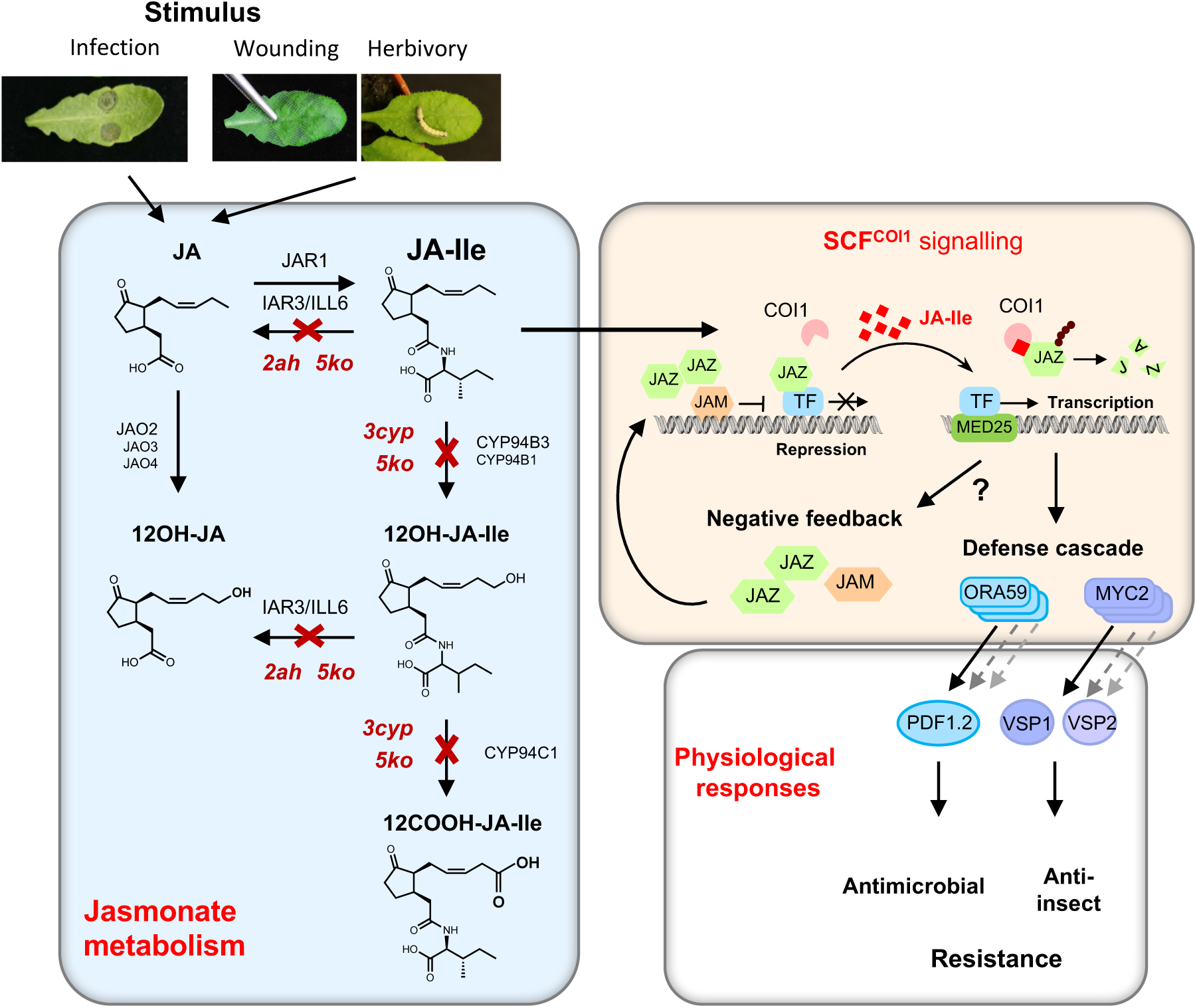
Positions of the impaired enzymatic steps in JA-Ile catabolic mutants and simplified view of analyzed signaling and defense responses. Left box: Necrotroph fungus infection or mechanical wounding/insect attack trigger jasmonic acid (JA) biosynthesis. Some JA is conjugated to isoleucine by JAR1 enzyme to form bioactive jasmonoyl-isoleucine (JA-Ile). JA-Ile is turned over by a two-step oxidation to 12OH-JA-Ile and 12COOH-JA-Ile by the cytochrome P450 enzymes CYP94B3/B1 and CYP94C1, or by conjugate cleavage mediated by the amidohydrolases (AH) IAR3 and ILL6. These AH also cleave 12OH-JA-Ile to release 12OH-JA. Red crosses indicate impaired JA-Ile catabolic steps in higher order mutants utilized in this study: oxidation-deficient (triple *cyp94b1 b3 c1* mutant, *3cyp*), deconjugation-deficient (double *iar3 ill6, 2ah*) or deficient for both pathways (quintuple mutant, *5ko*). Right box: in absence of JA-Ile, JAZ repressors block transcription of target genes. Upon biosynthesis, JA-Ile promotes the assembly of the COI1-JAZ co-receptor which is recruited into the SCF^COI1^ E3 ubiquitin ligase directing proteolytic degradation of JAZ repressors. Transcription factors (TFs) like MYC2 or ORA59 become activated in association with mediator 25 subunit (MED25) and allow the transcription of numerous stimulus-specific defense responses. The anti-insect branch was probed by the *MYC2-VSP* genes, and the antimicrobial branch by the *ORA59* and *PDF1.2* genes. Among JA-Ile targets are also genes encoding JAZ repressors or JAM proteins that compete with MYC2 and ORA59, to attenuate signaling. In mutant plants impaired for JA-Ile catabolism, overinduction of this negative feedback loop may occur and prevent hyper-stimulation of JA-Ile-dependent defenses and associated enhanced resistance.

Manipulating JA-Ile catabolic routes could be an attractive tool to engineer plants for enhanced pathogen resistance or modulate other JA-dependent processes. Loss- or gain-of-function mutant lines in *CYP94* or *AH* genes have revealed profound impacts on JA metabolism but contrasted consequences on JA responses. Arabidopsis lines ectopically overexpressing *CYP94* or *AH* generally have reduced JA-Ile levels and a metabolic shift towards oxidized and/or cleaved derivatives. As expected, increased turnover is correlated with attenuated JA responses, for example increased sensitivity to *Botrytis* infection or to insect larvae feeding (Aubert et al. 2015; Koo et al. 2011). In comparison, analysis of single, double, or triple mutant lines has not led to a unified conclusion as to how JA-Ile responses are impacted by gradual impairment of JA-Ile catabolism. Deficiency in CYP94 or AH expression impacts JA homeostasis consistently with their *in vitro* enzymatic activities. For example, *cyp94b3* or *b3c1* mutations lead to more JA-Ile upon wounding, concomitantly to reduced 12OH-JA-Ile and 12COOH-JA-Ile levels (Heitz et al. 2012; Kitaoka et al. 2011; Koo et al. 2011). Double *cyp94b3c1* and triple *cyp94b1b3c1* mutant plants exhibited slightly enhanced defense responses and tolerance to fungal infection (Aubert et al. 2015). Strikingly, in another study, the double *cyp94b1b3* and triple *cyp94b1b3c1* mutants exhibited deficient JA-Ile responses, questioning the current signaling model (Poudel et al. 2016). On the other hand, AH-deficient lines have not yet been tested for defense and resistance responses, so the contribution and impacts of this pathway are unknown.

To overcome gene redundancy and compensation mechanisms between pathways that occur frequently in JA metabolic mutants (Smirnova et al. 2017; Widemann et al. 2013), we introduced in the present study new plant lines with higher order mutations (Figure 1): a double *iar3 ill6* mutant (thereafter called *2ah*) impaired in the JA-Ile deconjugating pathway; and a quintuple *cyp94b1 cyp94b3 cyp94c1 iar3 ill6* (thereafter called *5ko*) deficient in all characterized enzymes turning over JA-Ile. We submitted these lines, along with the oxidation-deficient *cyp94b1b3c1* mutant (thereafter called *3cyp*), to parallel leaf stimulation by mechanical wounding or *B. cinerea* infection, two environmental cues triggering strong JA metabolism and signaling, but activating distinct transcriptional networks due to different cross-talks with other hormonal pathways (Pieterse et al. 2012). We determined the contribution of each catabolic pathway on JA-Ile turnover for each stress and investigated the impact of modified JA-Ile homeostasis on defense responses and tolerance to aggression by an herbivore insect or by fungal infection (Figure 1). The data highlight new stress-specific impacts of fully impairing either one or both JA-Ile catabolic pathways, where defense and aggressor resistance intensities are more correlated (negatively) to the hyper-activation of negative feedback signaling effectors rather than on actual JA-Ile levels.

## RESULTS

### Jasmonate profiling of higher order catabolic mutants reveals stimulus-specific impacts on hormone homeostasis

Effective inactivation of the respective genes was verified in each mutant line by RT-qPCR using RNA extracted from 1 h-wounded leaves (Supplemental Figure S1 and (Aubert et al. 2015)). The four plant genotypes in the Col-0 ecotype (WT, *2ah, 3cyp, 5ko*) were submitted separately to mechanical wounding or to *B. cinerea* inoculation. Mechanical wounding constitutes a synchronous and severe stimulus, and generates a massive jasmonate pulse where compound-specific dynamics can be followed (Chung et al. 2008; Glauser et al. 2008; Heitz et al. 2016). Therefore, a kinetic study was conducted with tissue collected at 1, 3 and 6 h post-wounding (hpw). Necrotic lesions inflicted by *B. cinerea* infection develop radially with fungal hyphae continuously infecting new tissue, and 3 days post-inoculation (dpi) constitutes an optimal time point for recording biochemical changes and for assessing antifungal resistance. We quantified levels of JA, 12OH-JA, JA-Ile and its catabolites 12OH-JA-Ile and 12COOH-JA-Ile in the two biological responses. JA levels were only marginally affected by the mutations (Supplemental Figure S2). Its oxidation product, 12OH-JA, is known to be partially formed via conjugate intermediates (Figure 1) upon wounding (Widemann et al. 2013) and accordingly, was less abundant in all mutants at 3 hpw but was only reduced in *2ah* at 6 hpw (Supplemental Figure S2). In contrast, 12OH-JA was less affected by mutations upon infection and was even enhanced in *5ko*, likely reflecting the predominant contribution of JAO enzymes directly oxidizing JA in this material (Figure 1) (Smirnova et al. 2017).

#### Impacts on JA-Ile homeostasis upon wounding

JA-Ile showed the typical pattern (Chung et al. 2008; Glauser et al. 2008; Heitz et al. 2012; Koo et al. 2011) in wounded WT leaves, peaking at 1 hpw and declined thereafter (Figure 2A). In *2ah* and *3cyp* lines, a significant overaccumulation of JA-Ile was recorded at 1 hpw and was prolonged in *3cyp*, extending data from mutants of lower order described previously (Heitz et al. 2012; Widemann et al. 2013; Zhang et al. 2016). In the *5ko* line that has both pathways impaired, a huge JA-Ile accumulation was recorded that culminated between 3-6 hpw close to 40-50 nmol g FW^-1^. These data show that in wounded leaves, both AH and CYP94 pathways contribute similarly to JA-Ile turnover and that their simultaneous inactivation synergistically boosts hormone hyperaccumulation at later time points. Profiles of CYP94-generated JA-Ile oxidation products were as expected: 12OH-JA-Ile levels were about double in *2ah* compared to WT, were suppressed in *3cyp* and of note, were higher in *5ko* than in *3cyp* at 6 hpw (Figure 2A). 12COOH-JA-Ile was only detected from 3 hpw and evolved similarly to 12OH-JA-Ile in *2ah* and *3cyp*, but was barely detected in *5ko*.

**Figure 2.**
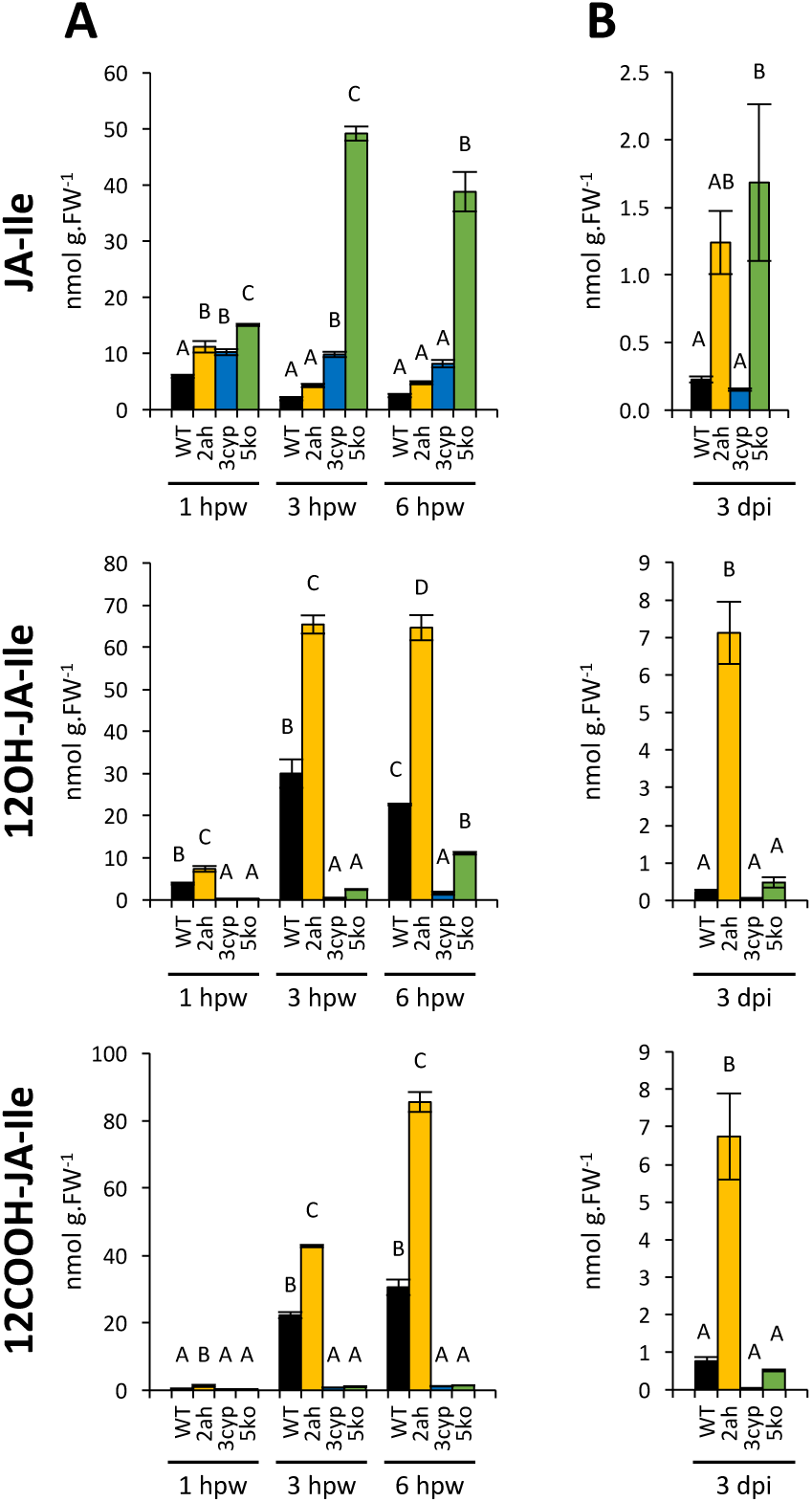
JA-Ile and oxidized catabolites accumulation in *2ah, 3cyp* and *5ko* mutants after mechanical wounding or *Botrytis cinerea* Infection. Six-week-old plants were wounded 3 times across leaf mid-vein (A) or drop-inoculated on two sites per leaf with a suspension containing 2.5×10^6^ fungal spores mL^-1^ (B). Treated leaves were harvested at 1, 3 and 6 hours post wounding (hpw) or 3 days post inoculation (dpi) on WT (black bars), *2ah* (yellow bars), *3cyp* (blue bars) and *5ko* (green bars) plants. Jasmonates were extracted and quantified by LC–MS/MS. JA-Ile, 12OH-JA-Ile and 12COOH-JA-Ile levels were expressed in nmol g^-1^ fresh weight (FW). Histograms represent the mean ± SEM of three (wounding) or four (*B. cinerea*) biological replicates. Columns labeled with different letters indicate a significant difference between genotypes at each time point as determined by one-way ANOVA with Tukey post-hoc test, P < 0.05.

#### Impacts on JA-Ile homeostasis upon Botrytis infection

The metabolic impacts were different in response to *B. cinerea* inoculation. JA-Ile levels were similar to WT in *3cyp*, confirming our previous data (Aubert et al. 2015) (Figure 2B). In contrast, JA-Ile levels were strongly enhanced in *2ah* but no further increase was observed in *5ko*. This result indicates that the amidohydrolase pathway is essential for JA-Ile clearance upon fungal infection and that blocking simultaneously the oxidative pathway does marginally impact JA-Ile accumulation. *2ah* also considerably enhanced 12OH-JA-Ile levels and 12COOH-JA-Ile, reinforcing initial data obtained with *iar3* or *ill6* single mutants (Widemann et al. 2013). In *2ah*, excess JA-Ile is likely oxidized into 12OH-JA-Ile that can no more be deconjugated by IAR3 and ILL6 and part of this enlarged pool is further oxidized to 12COOH-JA-Ile by CYP94C1. Together, these experiments demonstrate stimulus-specific contributions of AH and CYP94 catabolic pathways to JA-Ile turnover and accumulation of downstream derivatives.

### Impact of impaired catabolic pathways on defense and resistance responses is not reflecting JA-Ile accumulation in mutant plants

As the mutations altered the steady-state levels of bioactive JA-Ile, the lines were examined for induced expression of typical JA-regulated genes for each leaf stress model. In the case of wounding, transcripts of *MYC2*, an early-responsive transcription factor (TF) gene controlling late targets, accumulated similarly at 1 hpw, but declined less than WT in mutants at 3 hpw (*3cyp* and *5ko*) and 6 hpw (*2ah* and *5ko*) (Figure 3A). Two *VSP* genes were examined as late markers of the anti-insect branch of the JA defense pathway (Pieterse et al. 2012; Wasternack and Hause 2013). In WT, both *VSP1* and *VSP2* expression peaked at 3 hpw before declining to low levels at 6 hpw (Figure 3A). Expression was globally enhanced in mutant lines compared to WT, but with distinct patterns. *VSP1* was strongly enhanced in *2ah* at all time points, while in *3cyp* and *5ko*, gain in transcript levels was only observed at 6 hpw when WT signal faded. For *VSP2*, only *2ah* displayed enhanced expression at all 3 time points. This indicates that impaired JA-Ile catabolism results in persistent defense marker expression, but this is not commensurate with JA-Ile accumulation, because *5ko* displays lower defense than *2ah*. To determine the physiological impacts of such altered hormone and defense profiles, we used an insect feeding assay as a biological readout of signaling. When larvae of *Spodoptera littoralis* were placed on leaves of the four genotypes for 8 to 9 days, contrasted results were recorded: larvae fed on *2ah* plants were consistently lighter than those feeding on WT, indicating stronger anti-insect resistance (Figure 3B). This opposed to *3cyp*, that sustained higher insect development, confirming impaired resistance reported previously by Poudel et al. (2016). Finally, similarly to fungal infection, *5ko* displayed WT level of resistance to herbivory. These conclusions were drawn from 3 successive trials (Supplemental Figure S3) with large insect populations.

**Figure 3.**
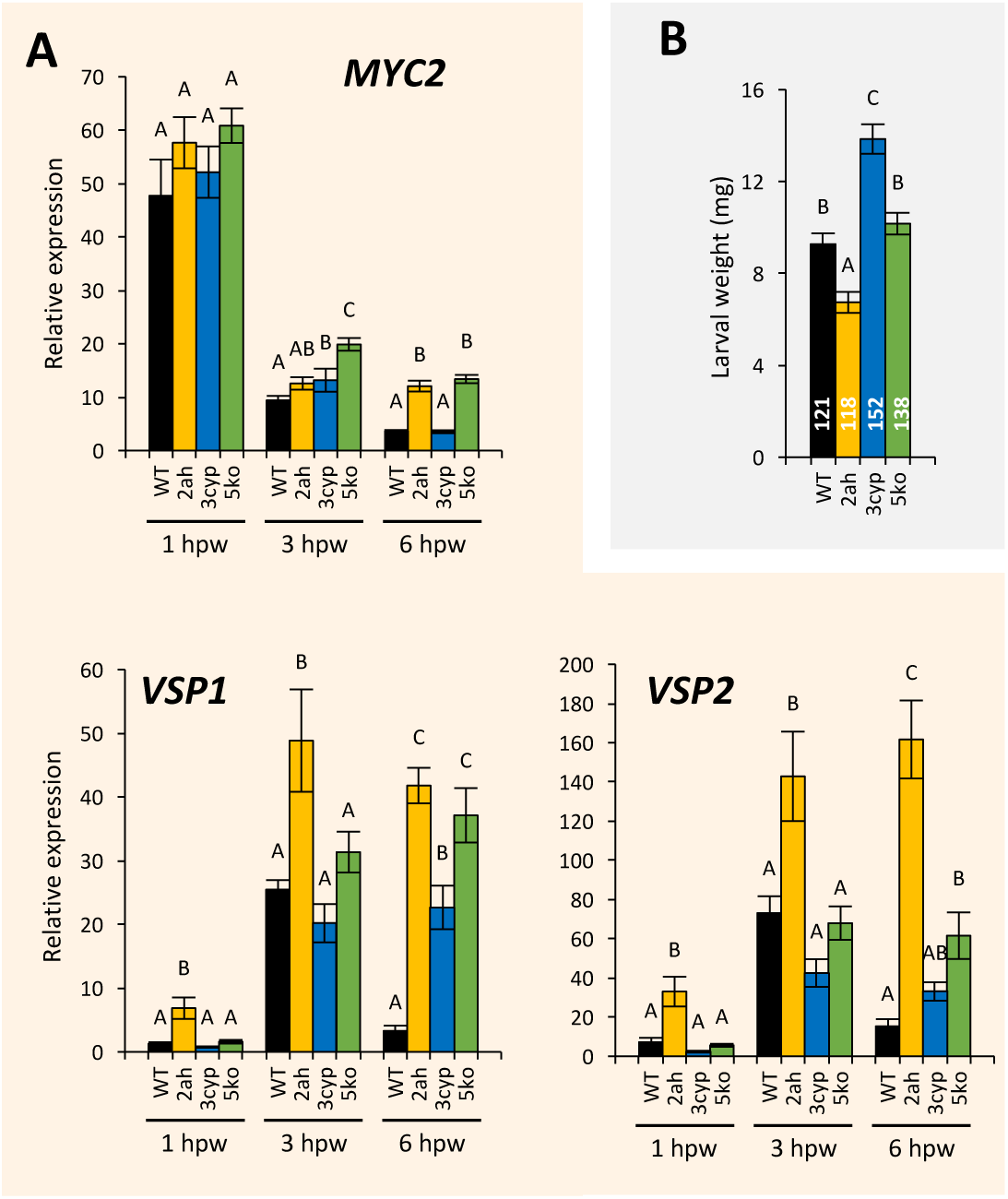
Expression profiles of JA-Ile-dependent genes in response to mechanical wounding and impact on insect feeding in JA-Ile catabolic mutants. (A) Expression profiles of jasmonate-dependent *MYC2, VSP1* and *VSP2* marker genes in response to wounding in WT (black bars), *2ah* (yellow bars), *3cyp* (blue bars) and *5ko* (green bars) mutants. Relative expression of each target gene at 1, 3 or 6 hours post wounding (hpw) is represented. Gene expression was determined by real-time PCR using gene-specific primers and normalized using *EXP* and *TIP41* reference genes. Transcript quantification was performed on three biological replicates analyzed in duplicate. Histograms represent mean expression ± SEM. Columns labeled with different letters indicate a significant difference between genotypes at each time point as determined by one-way ANOVA with Tukey posthoc test, P < 0.05. (B) Insect feeding assay. *S. littoralis* larvae were placed on leaves of 4-week-old plants. After 8-9 days, larval weight was determined. Histograms represent mean ± SEM from 3 independent trials (presented in Supplemental Figure S3). The number of total larvae on each genotype is indicated within the bars. Different letters indicate significant differences at *P* < 0.05 (linear mixed model).

A similar analysis was conducted for the response to *B. cinerea* infection. The antimicrobial branch of JA-Ile-dependent defense signaling was probed by monitoring *ORA59* TF and *PDF1.2* marker expression. Surprisingly, both genes exhibited reduced expression in *2ah* and *5ko* at 3 dpi, in contrast to *3cyp* that maintained WT levels (Figure 4A). Anti-fungal resistance assay indicated that only *3cyp* displayed smaller lesions, in agreement with our previous report (Aubert et al. 2015), while *3cyp* and to a lesser extent *2ah* supported reduced fungal biomass as estimated from *B. cinerea* cutinase signal (Figure 4B). Content in camalexin, a major antimicrobial phytoalexin in Arabidopsis, was determined but no significant differences were found between genotypes. Together, the data show that impairing either one or both JA-Ile catabolic pathways has distinct and stress-specific consequences on JA-mediated defense and resistance responses.

**Figure 4.**
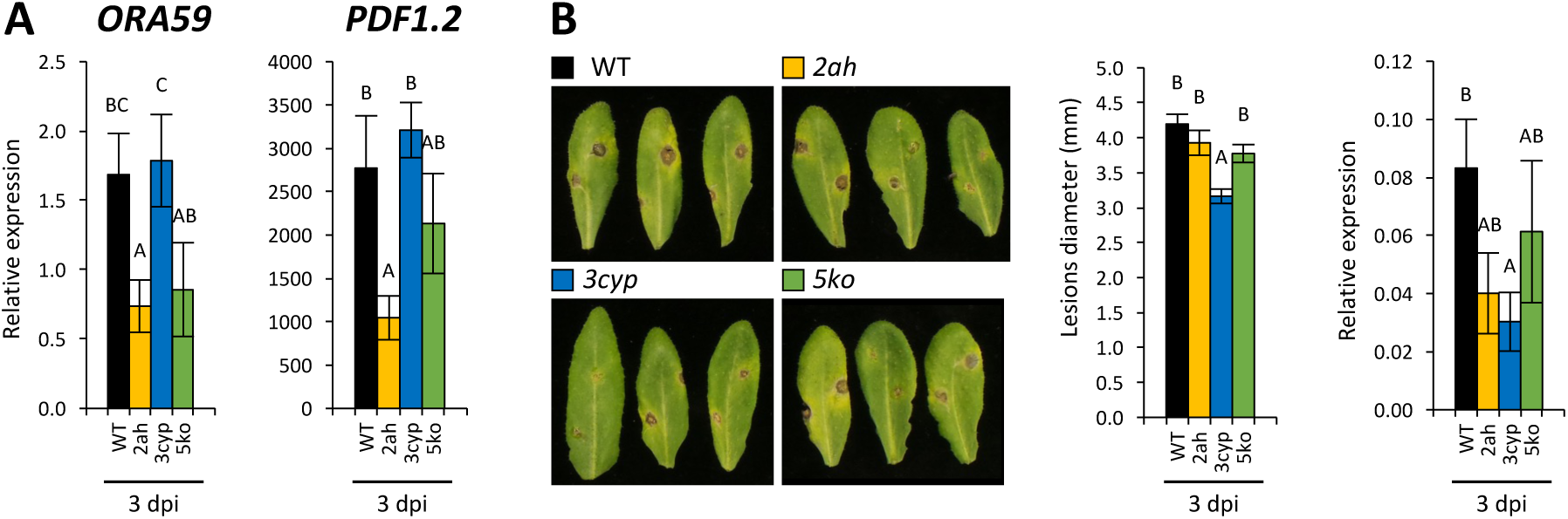
Expression profiles of JA-Ile-dependent defense genes and resistance levels in response to *B. cinerea* infection in *2ah, 3cyp* and *5ko* mutants. (A) Expression profiles of jasmonate-regulated *ORA59* and *PDF1.2* genes in response to infection by *B. cinerea* in WT (black bars), *2ah* (yellow bars), *3cyp* (blue bars) and *5ko* (green bars) mutants. Relative expression of each target gene at 3 day post infection (dpi) is represented. Gene expression was determined by real-time PCR using gene-specific primers and normalized using *EXP* and *TIP41* reference genes. Transcript quantification was performed on three biological replicates analyzed in duplicate. Histograms represent mean expression ± SEM. Columns labeled with different letters indicate a significant difference between genotypes at each time point as determined by one-way ANOVA with Tukey posthoc test, P < 0.05. (B) Disease resistance assessment. Two sites per leaf were inoculated across the main vein with 5 µL of spore suspension containing 2.5×10^6^ spores mL^-1^. Left panel: representative leaves of each genotype were detached and photographed at 3 dpi. Middle panel: histograms represent the mean lesion diameters ± SEM of about 100 lesion sites from 10 to 15 plants for each genotype. Right panel: evaluation of fungal growth by real-time qPCR with *B. cinerea* cutinase-specific primers on genomic DNA extracted from 3-day-infected leaves. Quantification was performed on three biological replicates analyzed in duplicate. Columns labeled with different letters indicate a significant difference as determined by one-way ANOVA with Tukey post-hoc test (P < 0.01).

### Impaired JA-Ile catabolism results in gain in resistance only when negative feedback effectors are not overinduced

Upon parallel investigation of the two leaf defense models, increased JA-Ile accumulation due to impaired catabolism did not correlate systematically with stronger defense or resistance phenotypes. Conversely, enhanced resistance can occur without elevated hormone levels (Figures 2, 3 and 4). These observations suggest that other players may be at work to limit or counteract overinduction of defense responses under deficient JA-Ile turnover. Obvious candidates for such negative regulation are genes encoding JAZ repressors (Browse 2009) or JAM bHLH factors competing with MYC transcription factors (Sasaki-Sekimoto et al. 2013), both classes of genes being JA-Ile-responsive. We selected *JAZ* genes that were previously described as highly induced and sensitive to catabolic pathway manipulation (Aubert et al. 2015; Heitz et al. 2012; Widemann et al. 2013). As shown in Figure 5A, *JAZ8, JAZ10, JAM1* and *JAM2* expression was maximal in all lines at 1 hpw with only slight enhancement in some mutants, but their transcripts were clearly more persistent in *3cyp* and *5ko* lines at 3 and 6 hpw, particularly for *JAZ*, compared to *2ah* and WT. This result establishes in *3cyp* and *5 ko* a correlation between hyperaccumulation of JA-Ile (Figure 2), the overinduction of *JAZ* and *JAM* genes (Figure 5), near-WT defense induction and WT (*5ko*) or deficient (*3cyp*) levels of insect resistance (Figure 3) compared to WT. In contrast, despite of enhanced JA-Ile levels upon wounding, *2ah* does not display persistent expression of these repressors (Figure 5, 3 and 6 hpw), and this correlates with more robust defense and reduced insect herbivory (Figure 3B). In response to *B. cinerea* infection, *JAZ1, JAZ8* and *JAM1* transcripts were overinduced in *2ah* and *5ko* (*JAM2* only in *5ko*) (Figure 5B), that overaccumulate JA-Ile, correlating with reduced defense (Figure 4A) and WT antifungal resistance (Figure 4B). On the contrary, these genes were not overinduced in *3cyp*, the only genotype exhibiting significantly increased antifungal resistance (Figure 4B). Therefore, the comparative analysis showed that elevated defense and resistance under deficient JA-Ile turnover is induced only if negative effectors are themselves not over-responding to JA-Ile accumulation.

**Figure 5.**
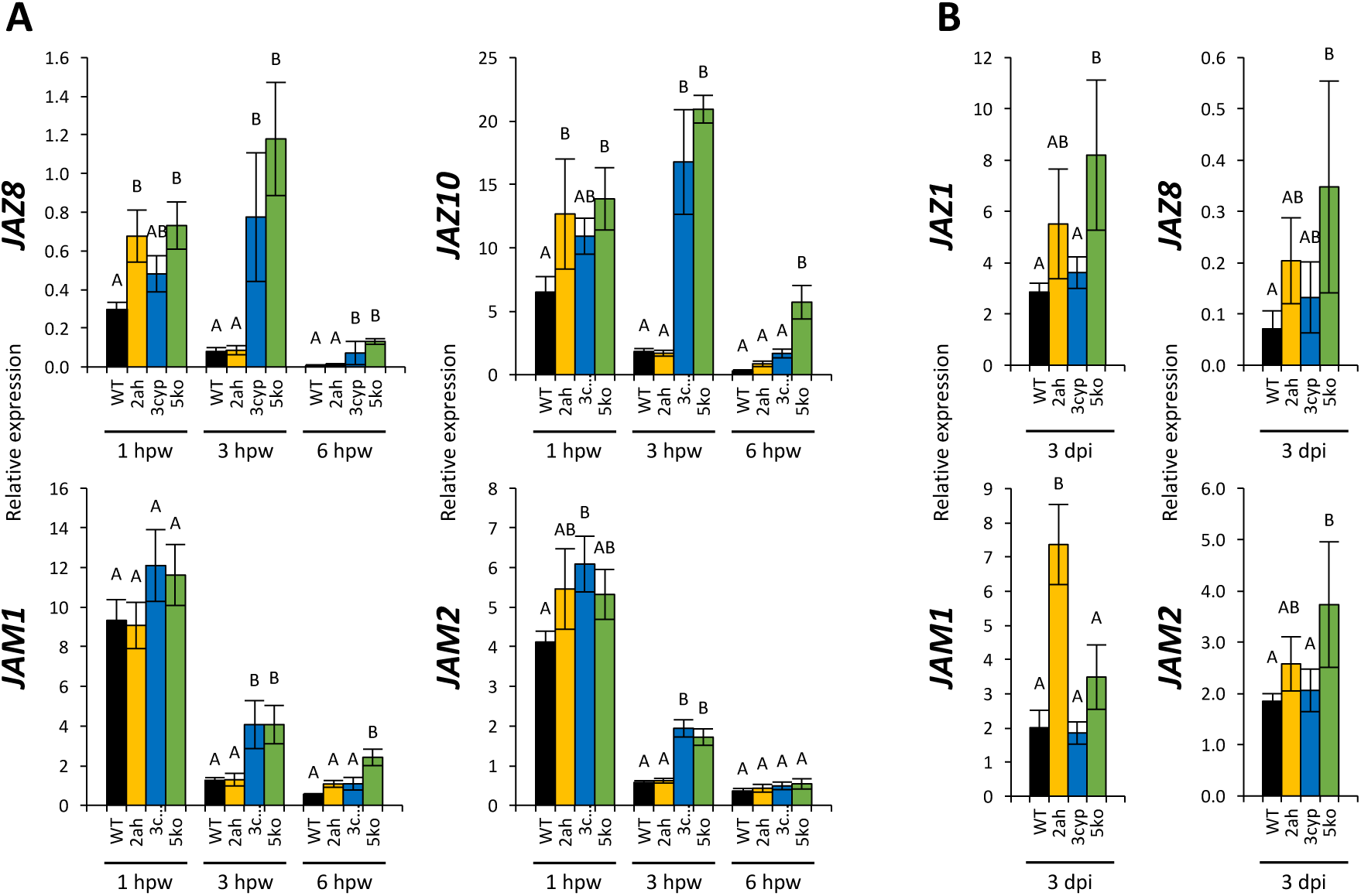
Expression profiles of JA-Ile dependant negative feedback genes in response to wounding and infection in *2ah, 3cyp* and *5ko* mutants. Expression profiles of jasmonate-dependent genes in response to wounding (A, *JAZ8, JAZ10, JAM1* and *JAM2*) or infection by *Botrytis cinerea* (B, *JAZ1, JAZ8, JAM1* and *JAM2*) in WT (black bars), *2ah* (yellow bars), *3cyp* (blue bars) and *5ko* (green bars) mutants. Relative expression of each target gene at 1, 3 or 6 hours post wounding (hpw) and 3 day post infection (dpi) is represented. Gene expression was determined by real-time PCR using gene-specific primers and normalized using *EXP* and *TIP41* reference genes. Transcript quantification was performed on three biological replicates analyzed in duplicate. Histograms represent mean expression ± SEM. Columns labeled with different letters indicate a significant difference between genotypes at each time point as determined by one-way ANOVA with Tukey posthoc test, P < 0.05.

**Figure 6.**
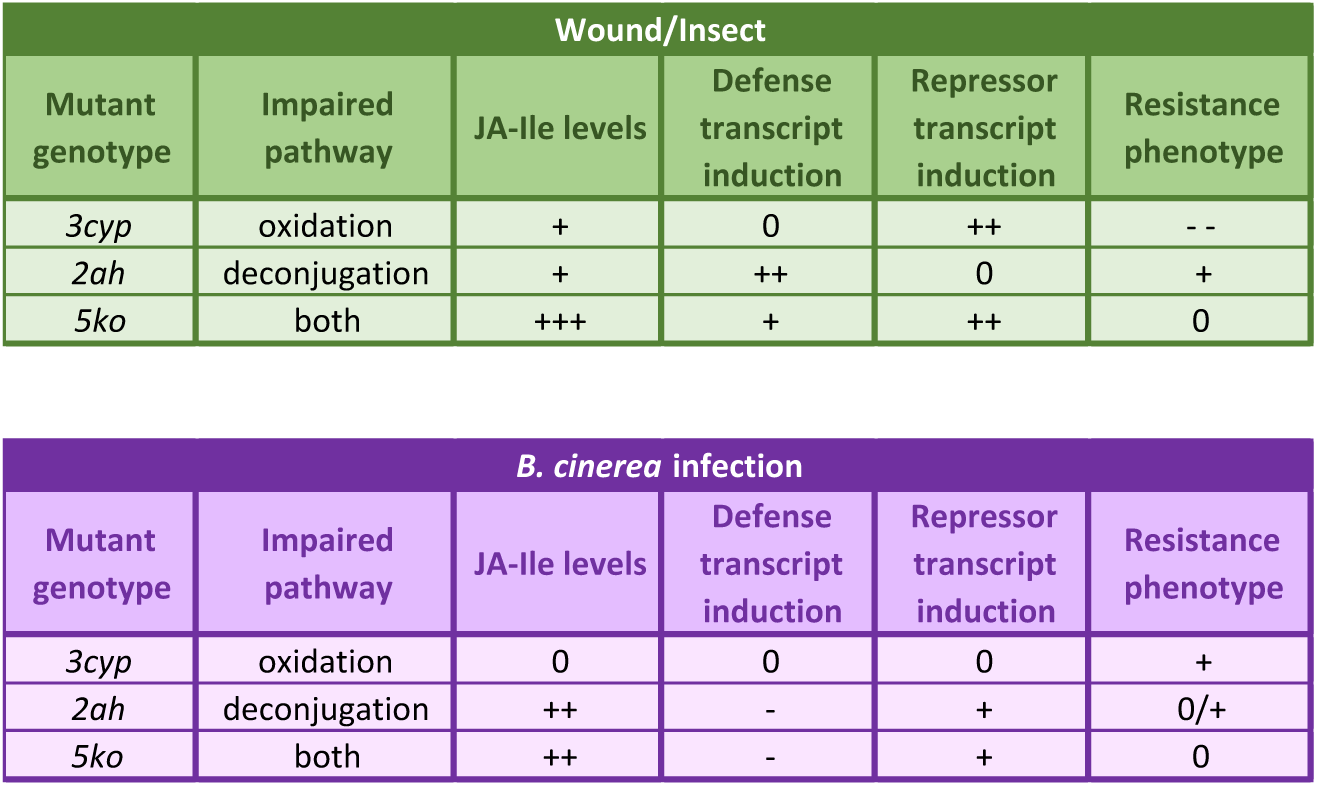
Relationships between impaired JA-Ile catabolism, JA-Ile levels, defense/repressor gene induction, and resistance to attackers. The data show distinct circuitry for the two leaf responses analyzed. The signs -, 0, or + indicate lower, equal, or increased response respectively, in stimulated mutant genotypes compared to WT. The two catabolic pathways have differential contributions to JA-Ile turnover in response to wounding or infection. JA-Ile accumulation is enhanced to different extents, except in infected *3cyp*, and this has variable consequences on defense amplitude and aggressor resistance. The trend emerging is that impaired hormone catabolism does, with or without JA-Ile overaccumulation, result in better defense/resistance, only if negative effectors like *JAZ* or *JAM* are themselves not overinduced.

To determine if JA-Ile catabolism mutations affected also basal gene expression in leaves before stress, we analyzed transcript levels of the same target genes in untouched leaves in a separate set of plants. Although higher fluctuations were recorded within a given genetic background than for stimulated tissues, two trends emerged: *MYC2* and *ORA59* TFs transcripts were significantly upregulated in *3cyp* and *5ko* lines, but levels of their respective targets *VSP* and *PDF1.2* were equal to or lower than WT in these lines (Supplemental Figure S4). Strikingly, all 5 analyzed genes encoding *JAZ* and *JAM* negative regulators were expressed at slightly but significantly higher levels in mutant (mostly *3cyp* and *5ko*) compared to WT. This result indicates that severe or even total block in JA-Ile catabolism does not increase basal expression of typical defense marker genes in Arabidopsis leaves.

## DISCUSSION

Hormonal compounds must be timely and spatially controlled to ensure coordinated physiological responses. Catabolic pathways have been characterized for all major plant receptor-binding hormonal ligands and are integral components of hormone homeostasis and action. Since their relatively recent discovery, JA-Ile catabolic pathways have been the focus of intense biochemical and physiological studies that disclosed their relationship with JA signaling. As a common theme in hormone catabolism (Kawai et al. 2014; Mizutani and Ohta 2010), an oxidative inactivation pathway was characterized, with CYP94 enzymes defining a two-step JA-Ile oxidation process. In contrast to most hormones for which conjugation corresponds to inactivation or generation of storage forms (Piotrowska and Bajguz 2011), JA requires conjugation to the amino acid isoleucine as the critical activation step. Consistently, deconjugation by the amidohydrolases IAR3 and ILL6 was characterized in Arabidopsis as a second JA-Ile removal pathway, acting in a JAR1 reverse reaction.

Understanding the functions of JA-Ile catabolism potentially offers avenues to tailor increased or on-demand defense signaling in diverse situations, and was addressed repetitively by loss-of-function studies in the two catabolic pathways (Aubert et al. 2015; Heitz et al. 2012; Kitaoka et al. 2011; Koo et al. 2011, 2014; Luo et al. 2016; Poudel et al. 2016; Woldemariam et al. 2012). In the present work, we conducted a comprehensive analysis of selected genotypes that allowed to evaluate the respective impact of each biochemical pathway on alterations in JA metabolism, defense regulation and resistance to biotic stress. The data clarify some former unexpected results and fill significant knowledge gaps by establishing novel pathway- and stress-specific features. It is remarkable that all mutant lines, including the newly generated *5ko* defective in all known JA-Ile-catabolizing enzymes, were undistinguishable from WT for growth under standard conditions. This comes along with the observation that CYP94 and IAR3/ILL6 deficiency does not increase basal JA-Ile accumulation or elevate defense gene expression in resting plants. We conclude that abolition of JA-Ile catabolism does neither turn on constitutive JA responses nor provoke growth inhibition.

Wounding/insect stress and infection by a necrotroph both induce strong JA metabolism and signaling (Campos et al. 2014), but due to different interactions with other hormonal networks (Pieterse et al. 2012), the transcriptome and defensive outcome display only partial overlap. Previous reports showed that single mutations in either the oxidative or cleavage pathway were sufficient to overaccumulate JA-Ile upon leaf wounding (Heitz et al. 2012; Kitaoka et al. 2011; Koo et al. 2011). This feature was extended in *3cyp* or *2ah* lines fully impaired in one or the other pathway, but here we demonstrate that their simultaneous deficiency raises JA-Ile to remarkably high levels, indicating that upon massive and rapid JA-Ile biosynthesis triggered by mechanical wounding, no additional enzymatic pathway can efficiently turnover and limit accumulation of JA-Ile. In response to *B. cinerea* infection, a comparable JA-Ile overaccumulation was recorded in *2ah* and *5ko* lines, but in marked contrast *3cyp* infected leaves accumulated WT levels of JA-Ile, suggesting that in this background, the activity of the AH pathway is sufficient to prevent abnormal hormone build-up. The CYP94 oxidation pathway is however active in infected WT plants, because oxidized derivatives are readily detected, albeit less abundantly than upon wounding. A preponderance of the AH pathway upon infection is supported by the strongly enhanced JA-Ile levels in *2ah* and *5ko* lines. This indicates that CYP94 cannot substitute for AH deficiency and that IAR3/ILL6 enzymes are essential for proper JA-Ile homeostasis upon necrotroph attack.

We next investigated the signaling output of perturbed JA-Ile homeostasis in the two leaf defense models and found that transcript levels of canonical regulatory and defense markers and resistance phenotypes were overall poorly correlated with changes in JA-Ile levels in mutant lines. Upon wounding, *3cyp* was closest to WT in term of *VSP* induction, contrasting with the deficient defense performance of this genotype. Both AH-deficient lines gained a more persistent *VSP1/2* expression, more pronounced for *VSP2* in *5ko*. This behavior was accompanied by a lower weight gain in *S. littoralis* larvae in *2ah*, but not in *5ko*, despite of its massive JA-Ile content. The situation with antimicrobial resistance also revealed a strong distortion of the usually linear relationship between hormone, defense and resistance levels, as the high JA-Ile-accumulating lines *2ah* and *5ko* exhibited depressed defense signaling and WT *B. cinerea* resistance. In contrast, *3cyp*, while maintaining WT JA-Ile levels, was able to better defend against the fungus, as reported earlier (Aubert et al. 2015). In this case, the normal behavior of the *ORA59-PDF1.2* module or camalexin levels do not seem to account for the better tolerance. Nevertheless, this result shows that more JA-dependent resistance can arise without an obvious over-accumulation of JA-Ile.

The JA-Ile regulatory network is complex and sets in motion cascades of positive and negative transcriptional regulators (Hickman et al. 2017). We hypothesized that known elements defining negative feedback loops may counteract some of the effects of chronic JA-Ile overaccumulation to restrain defense induction. This notion was introduced earlier for some *JAZ* genes in single or double mutants of one pathway, but its relevance was difficult to assess in single biological situations analyzed separately (Aubert et al. 2015; Heitz et al. 2012; Koo et al. 2011). Here we analyzed expression of *JAZ* and *JAM*, a second type of negative regulator inhibiting transcription of JA-Ile- and MYC-regulated target genes (Liu et al. 2019; Sasaki-Sekimoto et al. 2013) in the different pathway/genotype/stress combinations. We found a seemingly robust correlation between the overinduction of several *JAZ* and *JAM* genes and the absence of enhanced expression of defense response under impaired JA-Ile turnover (Figure 6). We selected *JAZ* genes that were previously found to be sensitive to altered JA-Ile homeostasis (Aubert et al. 2015; Heitz et al. 2012). *JAZ8, JAZ10, JAM1* and *JAM2* persisted longer than in WT in *3cyp* and *5ko* after wounding, but for unknown reasons behaved like WT in *2ah*, correlating with sustained *VSP* expression and better tolerance to feeding by *S. littoralis* larvae in this latter line. In the case of *B. cinerea* infection, a similar logic was respected, but in different genotypes. Both *2ah* and *5ko* displayed significantly higher expression of either *JAZ* or *JAM* or both genes, but this was not observed in *3cyp*. This correlated well with the former lines showing low defense and WT necrotic lesion sizes, and *3cyp* maintaining WT defense and being more resistant to infection. It should be noted that in *3cyp, ORA59-PDF1.2* as well as camalexin accumulation behaved like in WT, so other defense determinants must account for the observed phenotypes.

Together these results demonstrate that the two JA-Ile catabolic pathways contribute differentially to JA-Ile removal upon two distinct leaf stresses, and further variations may be expected in other organs or under other environmental constraints.

The impact of rewiring genetically JA-Ile catabolic capacity may vary between plant species and needs further investigations. For example, the oxidative or deconjugation pathway were silenced transiently (CYP94) or stably (JIH) in *Nicotiana attenuata*, and in two studies, clearly enhanced JA-Ile-dependent defenses and insect resistance were described (Luo et al. 2016; Woldemariam et al. 2012). We showed that it is possible to genetically extend the half-life and amplitude of JA-Ile burst in Arabidopsis, but this has contrasted consequences on signaling. The precise mechanism underlying control of signaling when JA-Ile catabolism is impaired remains unknown. Stress-specific “sensing” of excess JA-Ile on *JAZ* or *JAM* promoters may be mediated by the different interactions engaged between JA signaling and other hormonal pathways, such as the synergy with ethylene upon necrotroph attack, or with abscisic acid upon wounding or herbivory (Pieterse et al., 2012). The observation that blocking either catabolic pathway acting on the same substrate has distinct signaling and physiological outcomes suggests that signaling consequences of JA metabolism are not mediated solely through the control of JA-Ile levels. Modification of abundance of enzymatic products may also contribute to alter the signaling process(s). A recent paper reported that in addition to JA-Ile, 12OH-JA-Ile was an active jasmonate signaling through COI1 and contributing to wound responses (Poudel et al. 2019). Such an interpretation would be consistent with our description of metabolic and defensive features of *2ah* in wound/insect responses, but not with *3cyp* in the *Botrytis* assay. In addition, one must keep in mind that IAR3 also accepts auxin conjugates as substrates, which may also be at the basis of hormonal cross-talk and alter the signaling output (Widemann et al. 2013; Zhang et al. 2016). The only cases where restrained catabolism ends-up in enhanced tolerance to fungal or insect attack is when *JAZ* and/or *JAM* repressor transcripts are not overstimulated to reinforce negative feedback loops. The factors determining when negative regulators take control remain unknown but their elucidation will be of major importance to maximize the potential output of JA-regulated defenses. This provides the ground for future work at the protein, promoter and chromatin levels to determine how inhibitory mechanisms could be at work.

## METHODS

### Plant growth and treatments

*Arabidopsis thaliana* genotypes used were in the Col0 ecotype and were grown under a 12 h light/12 h dark photoperiod in a growth chamber. The individual T-DNA insertion lines were obtained from the Nottingham Arabidopsis Stock Center (NASC). The *3cyp* line was obtained by crossing the alleles *cyp94b1-1* (SALK_129672), *cyp94b3-1* (CS302217) and *cyp94c1-1* (SALK_55455) (Aubert et al. 2015). The *2ah* line was obtained after crossing the lines *iar3-5* (SALK_069047) and *ill6-2* (SALK_024894C). The quintuple *5ko* line was obtained by crossing the *3cyp* line with a double *iar3-5 ill6-1* (GK412-E11).

*B. cinerea* inoculation and resistance scoring were as described in (Aubert et al. 2015). For mechanical wounding experiments, 5 or 6 fully expanded leaves were wounded three times across mid-vein with a hemostat. At increasing time points following damage, leaf samples were quickly harvested and flash-frozen in liquid nitrogen before storing at −80°C until analysis.

### Insect performance assay

Four-week-old Arabidopsis plants were challenged with freshly hatched *Spodoptera littoralis* larvae (eggs obtained from Syngenta, Stein AG, Switzerland). Five larvae were placed on each of 11 pots, each containing 2 plants. Plants were placed in a transparent plastic box and kept in a growth chamber during the experiment (10 h light/14 h dark, 100 µmol m^-2^ s^-1^ of light, 20-22°C and 65% relative humidity). After 8-9 days of feeding, larvae were weighed on a precision balance (Mettler-Toledo, Greifensee, Switzerland). The experiment was performed successively three times (different sampling dates).

### RT-qPCR gene expression assays

Total RNA was extracted from plant leaves with TRI reagent (Molecular Research Center). One microgram of RNA was reverse transcribed using the SuperScript IV reverse transcription system (Thermo Fisher Scientific). Real-time PCR was performed on 10 ng of cDNA as described in Berr et al. (2010) using a LightCycler 480 II instrument (Roche Applied Science). The housekeeping genes EXP (At4g26410) and TIP41 (At4g34270) were used as internal references for qPCR on cDNA derived from infected/wounded or non-stimulated leaves. Measurement of fungal biomass was performed as described in (Smirnova et al. 2017) except that ACT2 (At3g18780) and UBQ10 (At4g05320) were used as reference genes. Gene-specific primer sequences used for qPCR are listed in Supplemental Table 1.

### Jasmonate and camalexin profiling

Jasmonates were identified and quantified by ultra high performance liquid chromatography coupled to tandem mass spectrometry (UHPLC-MS/MS). About 50-100 mg frozen plant material was extracted with 10 volumes (10 µL per mg) of ice-cold extraction solution (MeOH:water:acetic acid 70:29:0.5, v/v/v) containing 9,10-dihydro-JA and 9,10-dihydro-JA-Ile as internal standards for workup recovery. Grinding was performed with a glass-bead Precellys tissue homogenizer (Bertin Instruments, France) in 2 mL screw-capped tubes. After 30 min incubation at 4°C on a rotating wheel, homogenates were cleared by centrifugation before concentration of supernatants under a stream of N_2_ and overnight conservation at −20°C. After a second centrifugation step, extracts were submitted to quantitative LC-MS/MS analysis on an EvoQ Elite LC-TQ (Bruker) equiped with an electrospray ionisation source (ESI) and coupled to a Dionex UltiMate 3000 UHPLC system (Thermo). Five µL plant extract were injected. Chromatographic separation was achieved using an Acquity UPLC HSS T3 column (100 ×2.1 mm, 1.8 µm; Waters) and pre-column. The mobile phase consisted of (A) water and (B) methanol, both containing 0.1 % formic acid. The run started by 2 min of 95 % A, then a linear gradient was applied to reach 100 % B at 10 min, followed by isocratic run using B during 3 min. Return to initial conditions was achieved in 1 min, with a total run time of 15 min. The column was operated at 35 °C with a flow-rate of 0.30 mL min^-1^. Nitrogen was used as the drying and nebulizing gas. The nebulizer gas flow was set to 35 L h^-1^, and the desolvation gas flow to 30 L h^-1^. The interface temperature was set to 350 °C and the source temperature to 300 °C. The capillary voltage was set to 3.5 kV, the ionization was in positive or negative mode. Low mass and high mass resolution were 2 for the first mass analyzer and 2 for the second. Data acquisition was performed with the MS Workstation 8 for the mass spectrometry and the liquid chromatography was piloted with Bruker Compass Hystar 4.1 SR1 software. The data analysis was performed with the MS Data Review software. Absolute quantifications were achieved by comparison of sample signal with dose-response curves established with pure compounds and recovery correction based on internal standard signal. The transitions were, in negative mode: JA 209.3>59.3; JA-Ile 322.3>130.2; 12OH-JA-Ile 338.3>130.2; 12COOH-JA-Ile 352.2>130.1; 12OH-JA 225.2>59.3; in positive mode: camalexin 201.0>59.3.

### Statistical analysis

All statistical analysis was performed using InfoStat 2015d (http://www.infostat.com.ar). Comparisons of sample means were performed by one-way analysis of variance (*P* % 0.05 or *P* % 0.01) and Tukey’s post-hoc multiple comparisons tests (*P* < 0.05 or *P* < 0.01), and significant differences of means were determined.

## Supporting information

Marquis et al Supplemental data

## ACKNOWLEDGMENTS

J. Browse (WSU, Pullman, USA) for providing a segregating population of *iar3-5 ill6-2* line.

## REFERENCES

Aubert Y, Widemann E, Miesch L, Pinot F, Heitz T (2015) CYP94-mediated jasmonoylisoleucine hormone oxidation shapes jasmonate profiles and attenuates defence responses to Botrytis cinerea infection. J Exp Bot 66:3879–3892

Browse J (2009) Jasmonate passes muster: a receptor and targets for the defense hormone. Annu Rev Plant Biol 60:183–205

Bruckhoff V, Haroth S, Feussner K, Konig S, Brodhun F, Feussner I (2016) Functional Characterization of CYP94-Genes and Identification of a Novel Jasmonate Catabolite in Flowers. PLoS One 11:e0159875

Campos ML, Kang JH, Howe GA (2014) Jasmonate-triggered plant immunity. J Chem Ecol 40:657–675

Chini A, Fonseca S, Fernandez G, Adie B, Chico JM, Lorenzo O, Garcia-Casado G, Lopez-Vidriero I, Lozano FM, Ponce MR, Micol JL, Solano R (2007) The JAZ family of repressors is the missing link in jasmonate signalling. Nature 448:666–671

Chini A, Gimenez-Ibanez S, Goossens A, Solano R (2016) Redundancy and specificity in jasmonate signalling. Curr Opin Plant Biol 33:147–156

Chung HS, Koo AJ, Gao X, Jayanty S, Thines B, Jones AD, Howe GA (2008) Regulation and function of Arabidopsis JASMONATE ZIM-domain genes in response to wounding and herbivory. Plant Physiol 146:952–964

Gidda SK, Miersch O, Levitin A, Schmidt J, Wasternack C, Varin L (2003) Biochemical and molecular characterization of a hydroxyjasmonate sulfotransferase from Arabidopsis thaliana. J Biol Chem 278:17895–17900

Glauser G, Grata E, Dubugnon L, Rudaz S, Farmer EE, Wolfender JL (2008) Spatial and temporal dynamics of jasmonate synthesis and accumulation in Arabidopsis in response to wounding. J Biol Chem 283:16400–16407

Guo Q, Major IT, Howe GA (2018) Resolution of growth-defense conflict: mechanistic insights from jasmonate signaling. Curr Opin Plant Biol 44:72–81

Havko NE, Major IT, Jewell JB, Attaran E, Browse J, Howe GA (2016) Control of Carbon Assimilation and Partitioning by Jasmonate: An Accounting of Growth-Defense Tradeoffs. Plants (Basel) 5

Heitz T, Smirnova E, Widemann E, Aubert Y, Pinot F, Menard R (2016) The Rise and Fall of Jasmonate Biological Activities. Subcell Biochem 86:405–426

Heitz T, Widemann E, Lugan R, Miesch L, Ullmann P, Desaubry L, Holder E, Grausem B, Kandel S, Miesch M, Werck-Reichhart D, Pinot F (2012) Cytochromes P450 CYP94C1 and CYP94B3 catalyze two successive oxidation steps of plant hormone Jasmonoylisoleucine for catabolic turnover. J Biol Chem 287:6296–6306

Hickman R, Van Verk MC, Van Dijken AJH, Mendes MP, Vroegop-Vos IA, Caarls L, Steenbergen M, Van der Nagel I, Wesselink GJ, Jironkin A, Talbot A, Rhodes J, De Vries M, Schuurink RC, Denby K, Pieterse CMJ, Van Wees SCM (2017) Architecture and Dynamics of the Jasmonic Acid Gene Regulatory Network. Plant Cell 29:2086–2105

Kawai Y, Ono E, Mizutani M (2014) Evolution and diversity of the 2-oxoglutarate-dependent dioxygenase superfamily in plants. Plant J 78:328–343

Kitaoka N, Matsubara T, Sato M, Takahashi K, Wakuta S, Kawaide H, Matsui H, Nabeta K, Matsuura H (2011) Arabidopsis CYP94B3 encodes jasmonyl-L-isoleucine 12-hydroxylase, a key enzyme in the oxidative catabolism of jasmonate. Plant Cell Physiol 52:1757–1765

Koo AJ, Cooke TF, Howe GA (2011) Cytochrome P450 CYP94B3 mediates catabolism and inactivation of the plant hormone jasmonoyl-L-isoleucine. Proc Natl Acad Sci U S A 108:9298–9303

Koo AJ, Howe GA (2012) Catabolism and deactivation of the lipid-derived hormone jasmonoyl-isoleucine. Front Plant Sci 3:19

Koo AJ, Thireault C, Zemelis S, Poudel AN, Zhang T, Kitaoka N, Brandizzi F, Matsuura H, Howe GA (2014) Endoplasmic reticulum-associated inactivation of the hormone jasmonoyl-L-isoleucine by multiple members of the cytochrome P450 94 family in Arabidopsis. J Biol Chem 289:29728–29738

Liu Y, Du M, Deng L, Shen J, Fang M, Chen Q, Lu Y, Wang Q, Li C, Zhai Q (2019) MYC2 Regulates the Termination of Jasmonate Signaling via an Autoregulatory Negative Feedback Loop. Plant Cell 31:106–127

Luo J, Wei K, Wang S, Zhao W, Ma C, Hettenhausen C, Wu J, Cao G, Sun G, Baldwin IT, Wu J, Wang L (2016) COI1-Regulated Hydroxylation of Jasmonoyl-L-isoleucine Impairs Nicotiana attenuata’s Resistance to the Generalist Herbivore Spodoptera litura. J Agric Food Chem 64:2822–2831

Miersch O, Neumerkel J, Dippe M, Stenzel I, Wasternack C (2008) Hydroxylated jasmonates are commonly occurring metabolites of jasmonic acid and contribute to a partial switchoff in jasmonate signaling. New Phytol 177:114–127

Mizutani M, Ohta D (2010) Diversification of P450 genes during land plant evolution. Annu Rev Plant Biol 61:291–315

Pieterse CM, Van der Does D, Zamioudis C, Leon-Reyes A, Van Wees SC (2012) Hormonal modulation of plant immunity. Annu Rev Cell Dev Biol 28:489–521

Piotrowska A, Bajguz A (2011) Conjugates of abscisic acid, brassinosteroids, ethylene, gibberellins, and jasmonates. Phytochemistry 72:2097–2112

Poudel AN, Holtsclaw RE, Kimberlin A, Sen S, Zeng S, Joshi T, Lei Z, Sumner LW, Singh K, Matsuura H, Koo AJ (2019) 12-Hydroxy-jasmonoyl-L-isoleucine is an active jasmonate that signals through CORONATINE INSENSITIVE 1 and contributes to the wound response in Arabidopsis. Plant Cell Physiol pii: pcz109. doi:10.1093/pcp/pcz109

Poudel AN, Zhang T, Kwasniewski M, Nakabayashi R, Saito K, Koo AJ (2016) Mutations in jasmonoyl-L-isoleucine-12-hydroxylases suppress multiple JA-dependent wound responses in Arabidopsis thaliana. Biochim Biophys Acta 1861:1396–1408

Sasaki-Sekimoto Y, Jikumaru Y, Obayashi T, Saito H, Masuda S, Kamiya Y, Ohta H, Shirasu K (2013) Basic helix-loop-helix transcription factors JASMONATE-ASSOCIATED MYC2-LIKE1 (JAM1), JAM2, and JAM3 are negative regulators of jasmonate responses in Arabidopsis. Plant Physiol 163:291–304

Smirnova E, Marquis V, Poirier L, Aubert Y, Zumsteg J, Menard R, Miesch L, Heitz T (2017) Jasmonic Acid Oxidase 2 Hydroxylates Jasmonic Acid and Represses Basal Defense and Resistance Responses against Botrytis cinerea Infection. Mol Plant 10:1159–1173

Thines B, Katsir L, Melotto M, Niu Y, Mandaokar A, Liu G, Nomura K, He SY, Howe GA, Browse J (2007) JAZ repressor proteins are targets of the SCF(COI1) complex during jasmonate signalling. Nature 448:661–665

Wasternack C, Hause B (2013) Jasmonates: biosynthesis, perception, signal transduction and action in plant stress response, growth and development. An update to the 2007 review in Annals of Botany. Ann Bot 111:1021–1058

Wasternack C, Song S (2017) Jasmonates: biosynthesis, metabolism, and signaling by proteins activating and repressing transcription. J Exp Bot 68:1303–1321

Widemann E, Miesch L, Lugan R, Holder E, Heinrich C, Aubert Y, Miesch M, Pinot F, Heitz T (2013) The amidohydrolases IAR3 and ILL6 contribute to jasmonoyl-isoleucine hormone turnover and generate 12-hydroxyjasmonic acid upon wounding in Arabidopsis leaves. J Biol Chem 288:31701–31714

Widemann E, Smirnova E, Aubert Y, Miesch L, Heitz T (2016) Dynamics of Jasmonate Metabolism upon Flowering and across Leaf Stress Responses in Arabidopsis thaliana. Plants (Basel) 5

Woldemariam MG, Onkokesung N, Baldwin IT, Galis I (2012) Jasmonoyl-L-isoleucine hydrolase 1 (JIH1) regulates jasmonoyl-L-isoleucine levels and attenuates plant defenses against herbivores. Plant J 72:758–767

Yan C, Fan M, Yang M, Zhao J, Zhang W, Su Y, Xiao L, Deng H, Xie D (2018) Injury Activates Ca(2+)/Calmodulin-Dependent Phosphorylation of JAV1-JAZ8-WRKY51 Complex for Jasmonate Biosynthesis. Mol Cell 70:136–149.e7

Zhang T, Poudel AN, Jewell JB, Kitaoka N, Staswick P, Matsuura H, Koo AJ (2016) Hormone crosstalk in wound stress response: wound-inducible amidohydrolases can simultaneously regulate jasmonate and auxin homeostasis in Arabidopsis thaliana. J Exp Bot 67:2107–2120

