## Supplementary material for "Stress- and pathway-specific impacts of impaired jasmonoyl-isoleucine (JA-Ile) catabolism on defense signaling and biotic stress resistance in Arabidopsis": Marquis et al Supplemental data

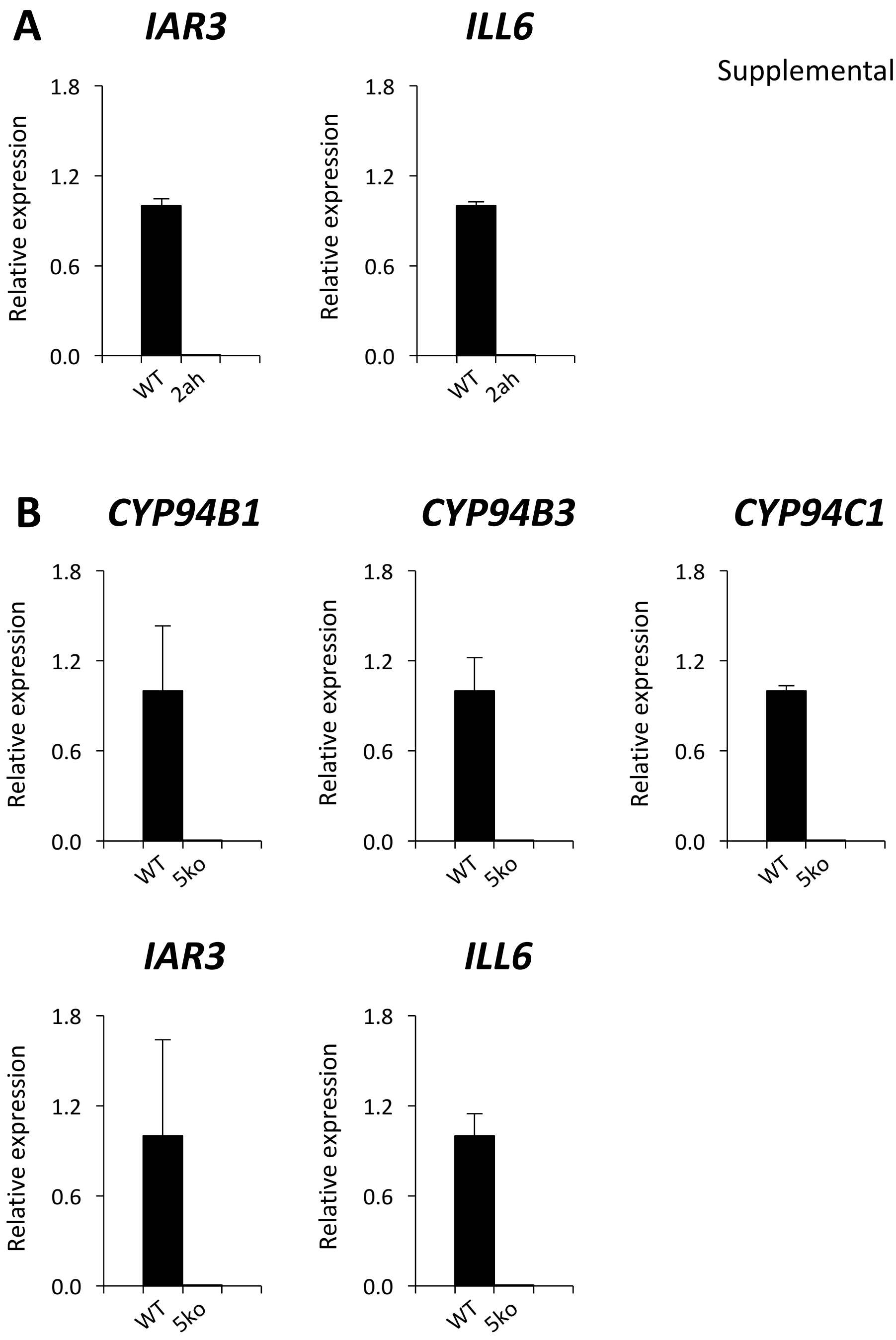


**Supplemental Figure S1. Characterization of *2ah* and *5ko* mutant lines used in this study.**

Expression of *CYP94B1*, *CYP94B3, CYP94C1, IAR3 and ILL6*, in WT, *2ah* (A) and *5ko* (B) mutant lines. Plants genotyped as homozygous were wounded and leaves were sampled after 1 h. RNA was extracted, and expression was analyzed by RT-qPCR using gene-specific primers and normalized using *EXP* and *GAPDH* reference genes. Expression in mutants is relative to expression in WT that was set to 1. The gene expression levels in three biological replicates were calculated using the $\Delta\Delta^{Ct}$ method Histograms represent mean expression of 3 biological replicates ± SD.

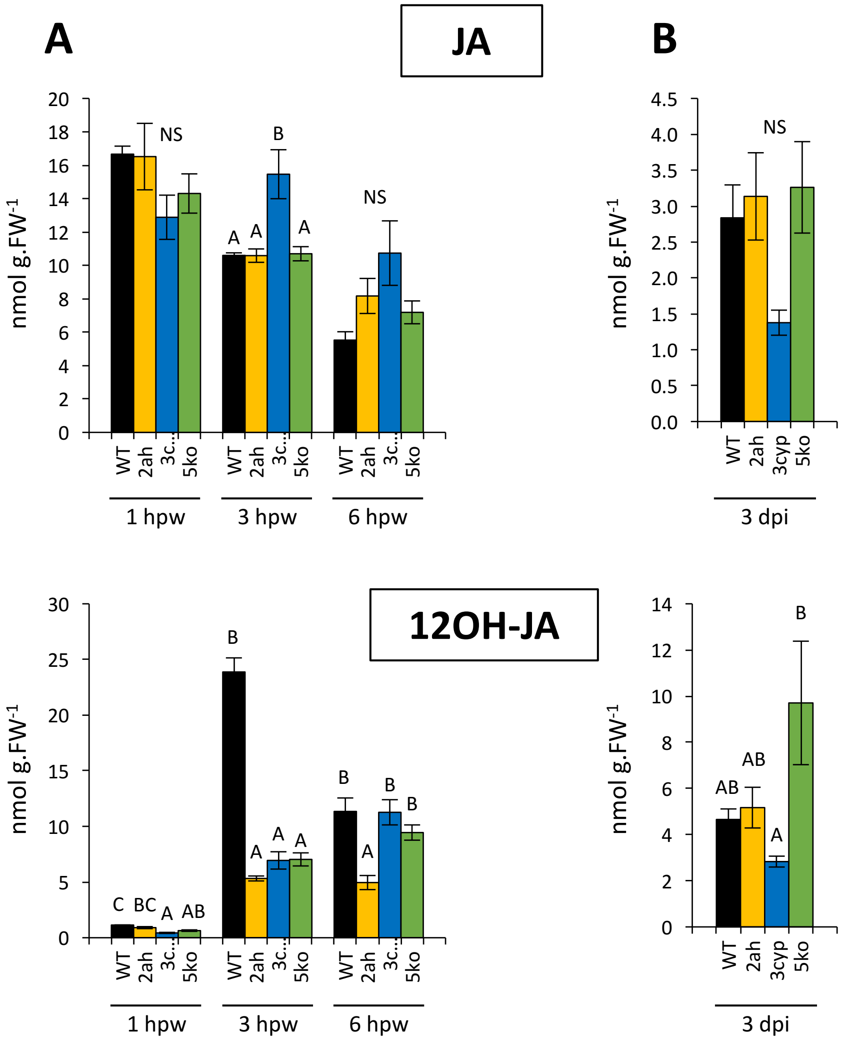


**Supplemental Figure S2. Jasmonate profiles in JA-Ile catabolic pathway mutants after wounding or *Botrytis cinerea* Infection.**

Six-week-old plants were either wounded (A) or drop-inoculated on two sites per leaf with a suspension containing 2.5x$10^6$ fungal spores $mL^{-1}$ (B). Leaves were harvested at 1, 3 and 6 h post wounding (hpw) or 3 days post inoculation (dpi) in WT (black bars), *2ah* (yellow bars), *3cyp* (blue bars) and *5ko* (green bars) mutants and jasmonates were extracted and quantified by LC–MS/MS. JA and 12OH-JA levels were expressed in nmol $g^{-1}$ fresh weight (FW). Histograms represent the mean ± SEM of three (wounding) or four (*B. cinerea*) biological replicates. Columns labeled with different letters indicate a significant difference between genotypes at each time point as determined by one-way ANOVA with Tukey post-hoc test, $P < 0.05$.

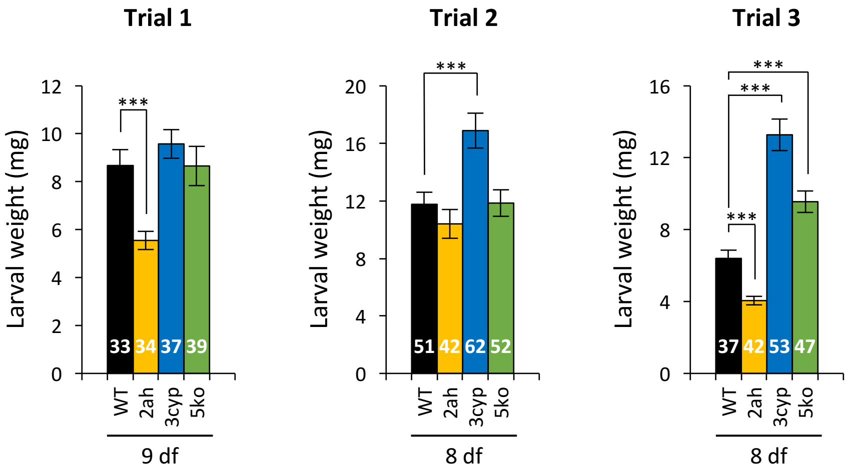


**Supplemental Figure S3. Susceptibility of WT, *2ah*, *3cyp* and *5ko* mutant lines to *Spodoptera littoralis* feeding.**

Freshly hatched *Spodoptera littoralis* larvae were placed on each genotype and larval weight was measured after 8 or 9 days of feeding (df) as indicated. Shown are mean ± SEM for 3 independent trials. The number of larvae measured for each population is indicated within the bars. Asterisks indicate a significant difference as determined by Student's *t* test (***P < 0.001)

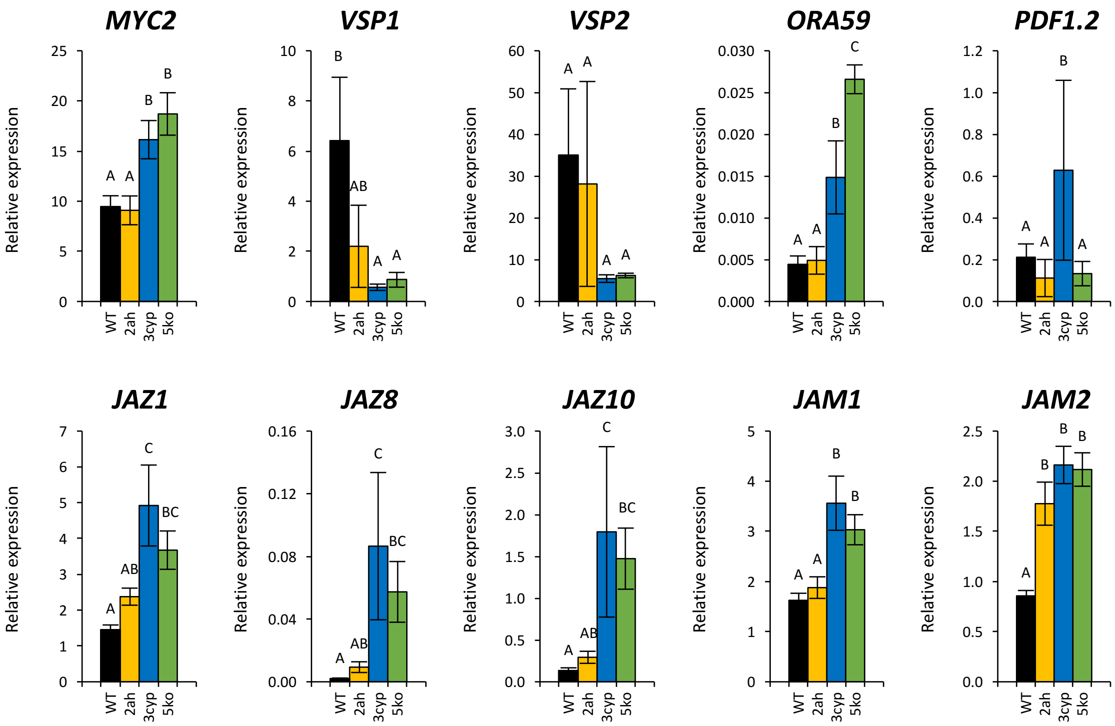


**Supplemental Figure S4. Basal expression of jasmonate-regulated defense genes in unstimulated leaves of WT (black bars), *2ah* (yellow bars), *3cyp* (blue bars) and *5ko* (green bars) lines.** Relative expression of each target gene determined by real-time qPCR using gene-specific primers and normalized using *EXP* and *TIP41* reference genes on three biological replicates analyzed in triplicate. Histograms represent mean expression ± SEM. Columns labeled with different letters indicate a significant difference between genotypes at each time point as determined by one-way ANOVA with Tukey post-hoc test, P < 0.01.

### Table S1. Primers used in the study

| **qPCR** | | | |
|---|---|---|---|
| **Gene/allele** | **Locus** | **Primer name** | **Sequence (5' -> 3')** |
| *EXP* | At4g26410 | At4g26410-qPCR_F | gagctgaagtggcttcaatgac |
| | | At4g26410-qPCR_R | ggtccgacatacccatgatcc |
| *TIP4.1* | At4g34270 | TIP41_LP | gtgaaaactgttggagagaagcaa |
| | | TIP41_RP | tcaactggataccctttcgca |
| *MYC2* | At1g32640 | E49-MYC2-L | gccgaaggaatacacgcaat |
| | | E50-MYC2-R | cgggttgtgaacgggcta |
| *VSP1* | At5g24780 | VSP1-F | ccgtcaatgtttggatctttg |
| | | VSP1-R | gctgtgttctcggtcccata |
| *VSP2* | At5g24770 | VSP2_qPCR_for | ggtgcccgcaaattgcaaagacta |
| | | VSP2_qPCR_rev | ggttgatgctccggtccctaacca |
| *ORA59* | At1g06160 | ORA59_qPCR_for_new | ttcgacgttgacatcttctcc |
| | | ORA59_qPCR_rev_new | tcttgcgtcataacaacactctg |
| *PDF1.2* | At5g44420 | E39-PDF1.2-L | caccccttatcttcgctgctctt |
| | | E40-PDF1.2-R | tacacttgtgtgctgggaagac |
| *JAZ1* | At1g19180 | JAZ1-F | ttctgagttcgtcggtagcc |
| | | JAZ1-R | cacgtctgtgagaagctaggc |
| *JAZ8* | At1g30135 | JAZ8-F | aatgtgttttcttcagatgttaccc |
| | | JAZ8-R | ttctctgcttgcgatcgatatt |
| *JAZ10* | At5g13220 | JAZ10-F | cgctcctaagcctaagttcca |
| | | JAZ10-R | tttcgaaatcgcaccttgaat |
| *JAM1* | At2g46510 | JAM1_qPCR_for | ggagctcacgcgtatcctaa |
| | | JAM1_qPCR_rev | ggattcgaagaagcagcaac |
| *JAM2* | At1g01260 | JAM2_qPCR_for | gaggccaatcaacgtgaagg |
| | | JAM2_qPCR_rev | tcccctcctccttctccatc |
| *CYP94B1* | At5g63450 | CYP94B1 qPCR F62 | caatgaggctttacccaccag |
| | | CYP94B1 qPCR R62 | aaatgtcgtcgtttgctgcat |
| *CYP94B3* | At3g48520 | CYP94B3 qPCR F62 | tggcttacacgaaggcttgtc |
| | | CYP94B3 qPCR R62 | agtcccacgaaactggaggat |
| *CYP94C1* | At2g27690 | CYP94C1 qPCR F63 | ggcccggattacgaagagttt |
| | | CYP94C1 qPCR R63 | ggccggaacttaccttcgtt |
| *IAR3* | At1g51760 | IAR3-0,4F | gttgctttaagggctgatatgg |
| | | IAR3-N69047-R | accgagaagcatcgtagtgtga |
| *ILL6* | At1g44350 | ILL6-GK212E12-F | gtgtcccatatccatccaacgg |
| | | ILL6-GK212E12-R | agactaatgaccgcggaagaag |
| *Actin2* | At3g18780 | Act_for | cttgcaccaagcagcatgaa |
| | | Act_rev | ccgatccagacactgtacttcctt |
| *UBQ10* | At4g05320 | UBQ10_for | ggccttgtataatccctgatgaataag |
| | | UBQ10_rev | aaagagataacaggaacggaaacatagt |
| *Cutinase* | B.c Z69264 | CutA-L | gatgtgacggtcatctttgccc |
| | | CutA-R | agatttgagagcggcgagg |
| **T-DNA genotyping** | | | |
| **Gene/allele** | | **Primer name** | **Sequence (5' -> 3')** |
| | | LBb1.3 (SALK) | attttgccgatttcggaac |
| | | GABI8409LB | atattgaccatcatactcattgc |
| *cyp94b1-1* | | 94B1-LP | tcgaatcacattgctctctcc |
| | | 94B1-RP | gggaattcactttcgaaatcc |
| *cyp94b3-1* | | CYP94B3-0,72F | gaacgtgggaagcgagaggaagc |
| | | BO1BG68 | tggtttggttctcactgttcac |
| *cyp94c1-1* | | SALK_055455 LP | tgtctttttggaaagtagcacc |
| | | SALK_055455 RP | gattccacggcctaaaagatc |
| *iar3-5 iar3-7* | | Salk_069047-LP | gttctccacgtgcgttatagc |
| | | Salk_069047-RP | aaaaagccacactgttccatg |
| *ill6-1* | | GK412E11-LP | gactatgcttcttggtgctgc |
| | | GK412E11-RP | cgcacctcttgaatacgtttc |
| *ill6-2* | | SALK_024894-LP | gactatgcttcttggtgctgc |
| | | SALK_024894-RP | cgcacctcttgaatacgtttc |
